# Identifying Key Cells for Fibrosis by Systematically Calling Cell Type–Phenotype Associations across Massive Heterogenous Datasets

**DOI:** 10.1101/2025.03.23.644786

**Authors:** Xi Chen, Yang Ding, Shiqi Yang, Lin Wei, Ming Chu, Hao Tang, Lei Kong, Yixin Zhou, Ge Gao

## Abstract

Fibrotic diseases pose a significant burden on health care, yet the key pathogenic fibroblasts involved remain unclear. We developed the fibrotic disease fibroblast atlas (FDFA), which comprises 394 single-cell and 38 spatial transcriptomic samples from 11 common fibrotic diseases.

To perform a cell-type phenotype association study in large-scale heterogeneous datasets, we developed the single-cell phenotype association research kit for large-scale dataset exploration (SPARKLE). SPARKLE handles heterogeneity by incorporating confounding information into its generalized linear mixed models (GLMMs).

The application of SPARKLE to FDFA revealed that matrix fibroblasts (MTFs) constitute a crucial pathogenic cell group in fibrosis. Their increased proportion correlate with the fibrotic process. MTFs also synergize with MYO-Fs, increasing their degree of fibrosis. Based on MTF, we identified 25 potential antifibrotic targets for broad-spectrum antifibrotic therapies.

This study enhances our understanding of fibrosis and provides a reliable framework for large-scale cell type–phenotype association research.

**Highlights:**

- A cross-tissue, multidisease fibrotic disease fibroblast atlas (FDFA) reveals diverse fibroblast subtypes in fibrosis diseases.
- A new toolkit, SPARKLE, is introduced for detecting robust cell type–phenotype associations in large-scale heterogeneous datasets.
- Multiple pathogenic fibroblasts associated with common fibrosis disease (MTF) onset and progression have been identified, suggesting potential cell-specific therapeutic strategies for fibrosis-related diseases.

## Introduction

Fibrosis is emerging as a significant global health crisis, affecting 25% of the global population^1,2^. With no effective drug treatments available, fibrotic diseases are considered irreversible and fatal, contributing to ∼45% of deaths in developed nations^3,4^. These diseases impact nearly all major tissues, including idiopathic pulmonary fibrosis (IPF), ischemia cardiomyopathy (ICM), viral liver cirrhosis (VLC), chronic kidney disease (CKD), arthrofibrosis (AF), and hypertrophic skin scarring (HSS)^5^. However, the common pathological mechanisms underlying fibrosis remain largely elusive, especially whether there is a set of common “pathogenic cells” that contribute to the onset and progression of fibrotic diseases^6,7^.

Here, we integrated single-cell transcriptome data from 394 samples and spatial transcriptome data from 38 samples across 11 common fibrotic diseases to establish the first human cross-tissue, multidisease fibrotic disease fibroblast atlas (FDFA). In large-scale single-cell studies, data heterogeneity caused by multiple intrinsic and extrinsic factors is quite common^8–10^. To address heterogeneity in cell proportion–phenotype association analysis, we developed a computational toolkit based on generalized linear mixed models (GLMMs): the single-cell phenotype association research kit for large-scale dataset exploration (SPARKLE). Using SPARKLE, we identified matrix fibroblasts (MTFs) as a key pathogenic cell group that promotes fibrosis via three distinct mechanisms. Notably, SPARKLE helped identify 25 potential antifibrotic therapeutic targets and five effective broad-spectrum antifibrotic drugs targeting MTF, suggesting a promising clinically efficient approach for drug repurposing.

## Results

### Construction of a cross-tissue fibrotic disease fibroblast atlas (FDFA)

To construct a comprehensive human cross-tissue, multidisease fibrotic disease fibroblast atlas (FDFA), we curated 6,757 publicly available single-cell datasets (34,589 samples) published between February 2009 and August 2023. We initially removed 2,940 samples with incomplete information, resulting in 31,649 samples (Fig. 1A). Of these, 15,848 nonhuman samples were excluded, leaving 15,801 human single-cell samples. Using ScType for automated annotation, we excluded 11,510 samples that did not contain fibroblasts, retaining 4,291 samples with fibroblasts, totaling approximately 1.9 million fibroblasts. The metadata are further screened to exclude nonfibrosis-related diseases while normal fetal and adult samples are retained as controls. Finally, we compiled 432 samples for 11 common fibrotic diseases across 6 tissue types, a total of 147,254 fibroblasts from 394 single-cell transcriptomic samples and 38 spatial transcriptomic samples, to construct the fibrotic disease fibroblast atlas (FDFA, Fig. 1A and Supplementary Fig. 1A-1F).

**Figure 1.**
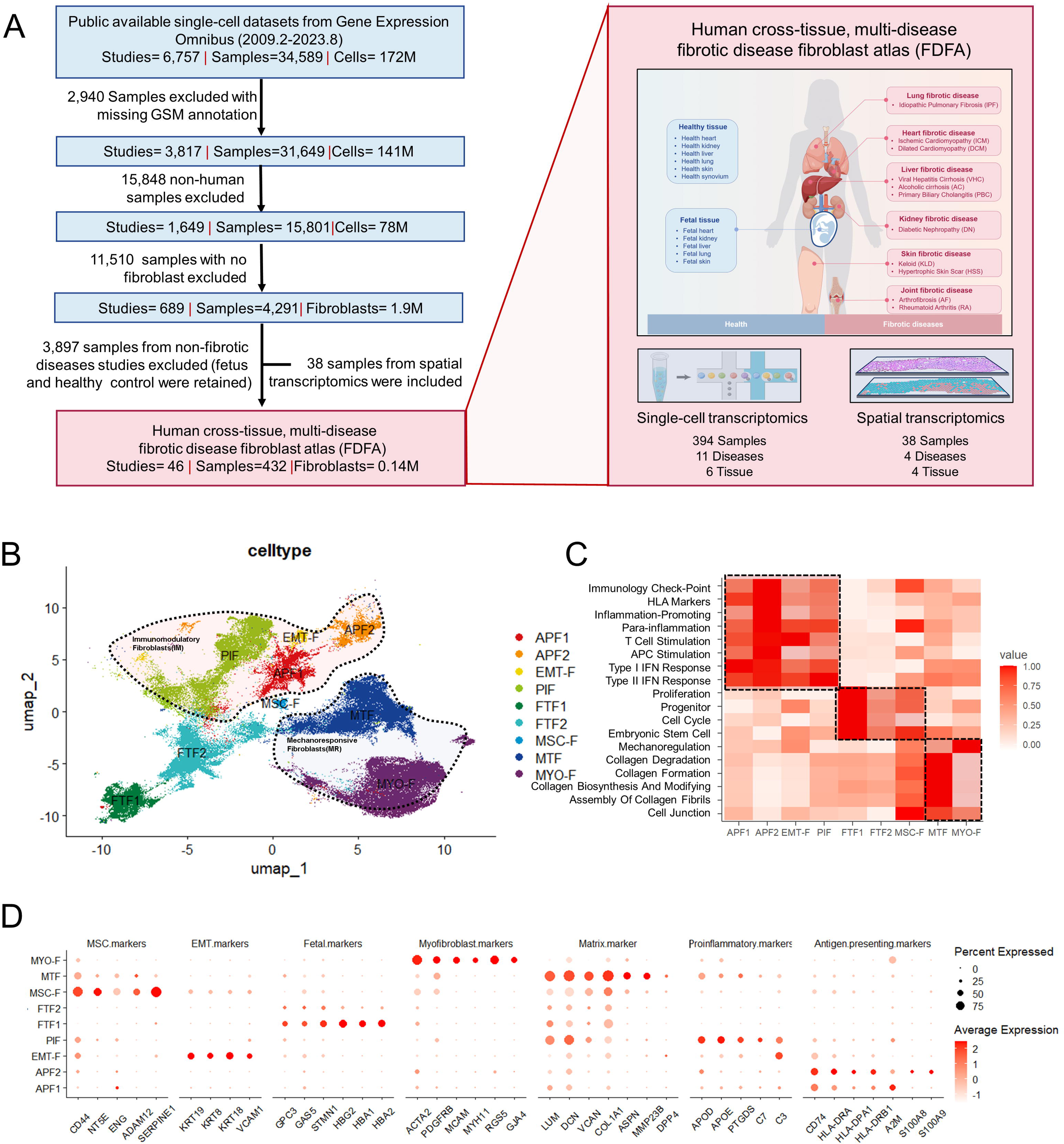
Construction and Analysis of the Fibrotic Disease-Related Fibroblast Atlas. (A) Schematic diagram of the fibrotic diseases fibroblast atlas (FDFA) construction. This atlas integrates single-cell transcriptomics and spatial transcriptomics data from various fibrotic diseases across different tissues. Single-cell transcriptomics includes data from 147,254 fibroblasts, 394 samples, 11 diseases, and 6 tissues. Spatial transcriptomics includes data from 38 samples, 4 diseases, and 4 tissues, enabling comprehensive cell type annotation, phenotype association analysis, and multimodal intersection analysis. (B) UMAP visualization of fibroblasts across different tissues. Each color represents a different fibroblast subtype. The dashed lines highlight superclusters with similar functional annotations, indicating potential functional similarities among these subtypes. (C) Heatmap showing the functional analysis of nine different cell types. The values in the heatmap represent the functional scores of specific gene sets related to various cellular functions. The dashed lines indicate superclusters of fibroblast subtypes with shared functional characteristics. (D) Dot plot of different cell marker genes. The size of each dot represents the percentage of cells expressing the gene within a cluster, whereas the color intensity represents the expression level of the gene. This visualization aids in identifying the distribution and expression levels of key marker genes across different fibroblast subtypes.

Based on the FDFA, three distinct superclusters were identified, comprising nine novel subtypes (Fig. 1B). The naïve fibroblast supercluster is composed mainly of fetal-derived fibroblasts, including fetal fibroblast type 1 (FTF1), fetal fibroblast type 2 (FTF2), and mesenchymal stem cell-like fibroblasts (MSC-F). FTF1 and FTF2 express fetal markers such as GPC3 and STMN1, which are associated with the cell cycle and proliferation (Fig. 1C). MSC-like fibroblasts express markers such as CD44, NT5E, and ENG and are present in all tissues except skin, with the highest concentrations in synovial tissue and lungs (Supplementary Fig. 1G-1H).

The immunomodulatory fibroblast supercluster includes antigen-presenting fibroblasts (APF1 and APF2), proinflammatory fibroblasts (PIF), and epithelial–mesenchymal transition fibroblasts (EMT-F) (Fig. 1B). APF1 and APF2 express antigen-presenting markers such as CD74 and HLA-DPA1, which play critical roles in T-cell activation (Fig. 1D). PIFs express proinflammatory markers such as APOD and APOE, which facilitate inflammatory responses. EMT-F involves the expression of epithelial markers such as KRT19 and KRT8, potentially representing an intermediate stage in the transition from epithelial cells to myofibroblasts.

The mechanoresponsive fibroblast supercluster consists of matrix fibroblasts (MTFs) and myofibroblasts (MYO-Fs) (Fig. 1B). MTFs, characterized by high expression of collagen and ECM-related genes, are vital for ECM synthesis and organization (Fig. 1C). MYO-Fs, which are marked by ACTA2 and PDGFR, play significant roles in wound healing and have contractile properties that aid in wound contraction (Fig. 1D).

### Development of a model-based toolkit for detecting cell type–phenotype associations across large-scale heterogeneous datasets

When performing joint analyses of single-cell data from different sources, significant heterogeneity presents a considerable challenge, especially for detecting robust cell type–phenotype associations^11^. Multiple intrinsic and extrinsic factors during study design and execution contribute to heterogeneity: differences in population structure and sampling methods can lead to severe selection bias^12^; variations in sequencing platforms and data analysis techniques may introduce information bias^13^ (Figure 2A).

**Figure 2.**
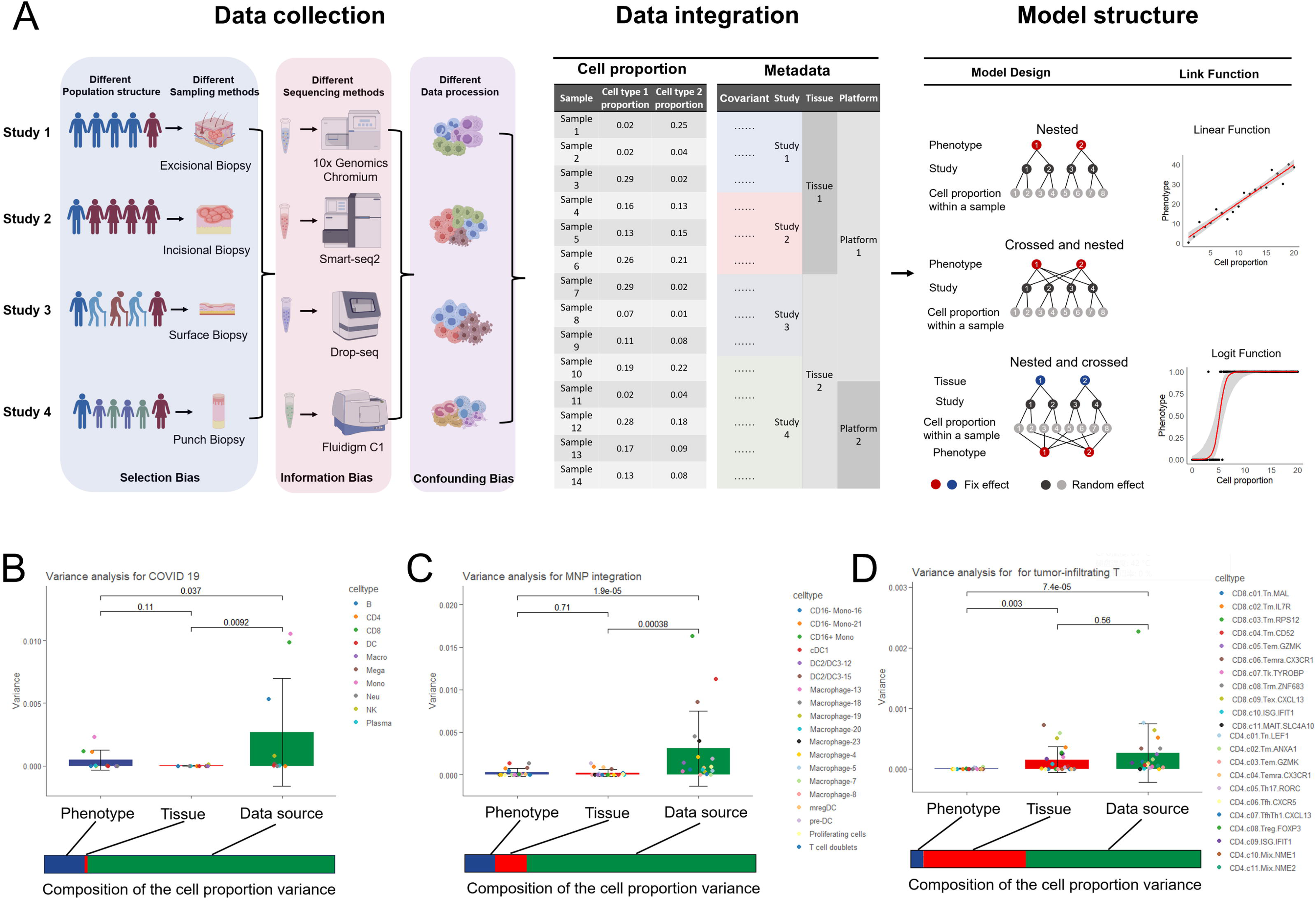
SPARKLE Architecture and Its Significance in Data Integration. (A) Schematic representation of the SPARKLE computational workflow. (B-E) Variance analysis results of the associations between phenotypes, tissues, data sources, and cell proportions across different datasets. Each colored dot represents the variance in cell proportions for different cell types within each grouping variable. The color bar under each plot indicates the composition of the cell proportion variance. (B) The COVID-19 dataset. (C) The human monocyte and macrophage dataset. (D) The tumor-infiltrating T-cell dataset.

While plain statistical test-based methods such as the Wilcoxon rank-sum test have been widely used^14–16^, they often fail to handle heterogeneity robustly. To address this issue, we selected three representative large-scale atlas datasets with complete metadata: the COVID-19 atlas, the tumor-infiltrating T-cell atlas, and the human monocyte and macrophage atlas (MNP atlas)^17–19^. The COVID-19 dataset contains 249 samples from the same tissue source (PBMC) but collected from 12 different cities, resulting in 17 distinct batches. In contrast, samples in the tumor-infiltrating T-cell atlas and the human monocyte and macrophage atlas are derived from various tissue sources. The human monocyte and macrophage atlas includes 178,651 cells from 13 tissue types, whereas the tumor-infiltrating T-cell atlas comprises 390,000 cells from 21 cancer types. Notably, the greater diversity of data sources in the tumor-infiltrating T-cell atlas suggests a greater potential for heterogeneity than does the MNP atlas. Further inspection revealed that in the tumor-infiltrating T-cell atlas, removing only one study^20^ led to changes in up to 30.4% (7 out of 23) of the cell type–phenotype conclusions identified by the Wilcoxon test (Supplementary Table 2). Similarly, in the monocyte atlas, removing a single study^21^ resulted in changes up to 26.3% (5 out of 19) of the cell type conclusions. Even the removal of just one batch in the COVID-19 dataset caused a shift in up to 30% (3 out of 10) of the cell type-phenotype conclusions based on the Wilcoxon test (Supplementary Table 2).

Notably, variance analysis demonstrated that across all datasets, the variance stemming from the data source was significantly greater than that stemming from phenotype differences (Figure 2B-D), suggesting that incorporating covariate information into cell type–phenotype associations is crucial and can potentially mitigate the impact of single-cell data heterogeneity on the final results. Thus, we constructed four linear regression models to assess how different covariates influence association analysis outcomes. In the absence of covariates (Model 1), the p values for cell type–phenotype associations closely resembled those from the Wilcoxon test. However, adding tissue information (Model 2), data source grouping (Model 3), or both (Model 4) significantly altered the p values, affecting the statistical significance (p < 0.05) of these associations (Supplementary Fig. 2A-2C). When the Wilcoxon test results without any covariates were used as a reference, we observed that more than 30% of the cell type–phenotype association conclusions in most datasets (and even more than 50% in some datasets) changed upon adding covariates (Supplementary Fig. 3A-3C). These results highlight the importance of including covariates in large-scale data integration analyses, even basic ones such as tissue type and data source.

To detect phenotype associations across large-scale single-cell datasets robustly, we developed the single-cell phenotype association research kit for large-scale exploration (SPARKLE, Fig. 2A and Supplementary Fig. S8). Briefly, SPARKLE first performs a Hausman test to determine whether covariates should be modeled as fixed or random effects, depending on the type of covariates. For random-effect covariates, SPARKLE supports comprehensive modeling of nested, crossed and nested, and nested and crossed structures (Fig. 2A and Supplementary Fig. S8), along with multiple link functions, including logit and linear functions. Notably, bootstrap procedures suggested that the output of SPARKLE is rather robust: almost all cell type–phenotype associations identified by SPARKLE showed high (>80%) consistency among different setups (while not for the canonical Wilcoxon-based approach, Supplementary Fig. 4A-4F).

### Exploration of cell type–phenotype associations in the fibrotic disease fibroblast atlas with SPARKLE

In an attempt to identify fibrosis-associated fibroblasts, we ran SPARKLE on a fibrotic disease fibroblast atlas (Fig. 3A-3B; also see Methods and Supplementary Figs. 5A-5B, 6A-8B, and 7A-6B for more details). The output of SPARKLE clearly revealed that MTF was significantly positively correlated with fibrosis phenotype, whereas APF1, FTF1 and PIF were significantly negatively correlated (Fig. 3B). Interestingly, the results identified MYO-F as a positively correlated factor, but only when the MTF was modeled as a significant confounding factor (Fig. 3C). To elucidate the complex relationships between these cell types, we further used mediation analysis to explain the observed phenomena (Fig. 3D). The mediating effects of PIF, FTF1 and APF1 had the same sign as the main effect, indicating that MTF played an indirect mediating role in the association between PIF and the phenotype. In contrast, the mediation effects of MYO-F had opposite signs to the main effects, indicating that MTF had a masking effect on the association between MYO-F and the phenotype.

**Figure 3:**
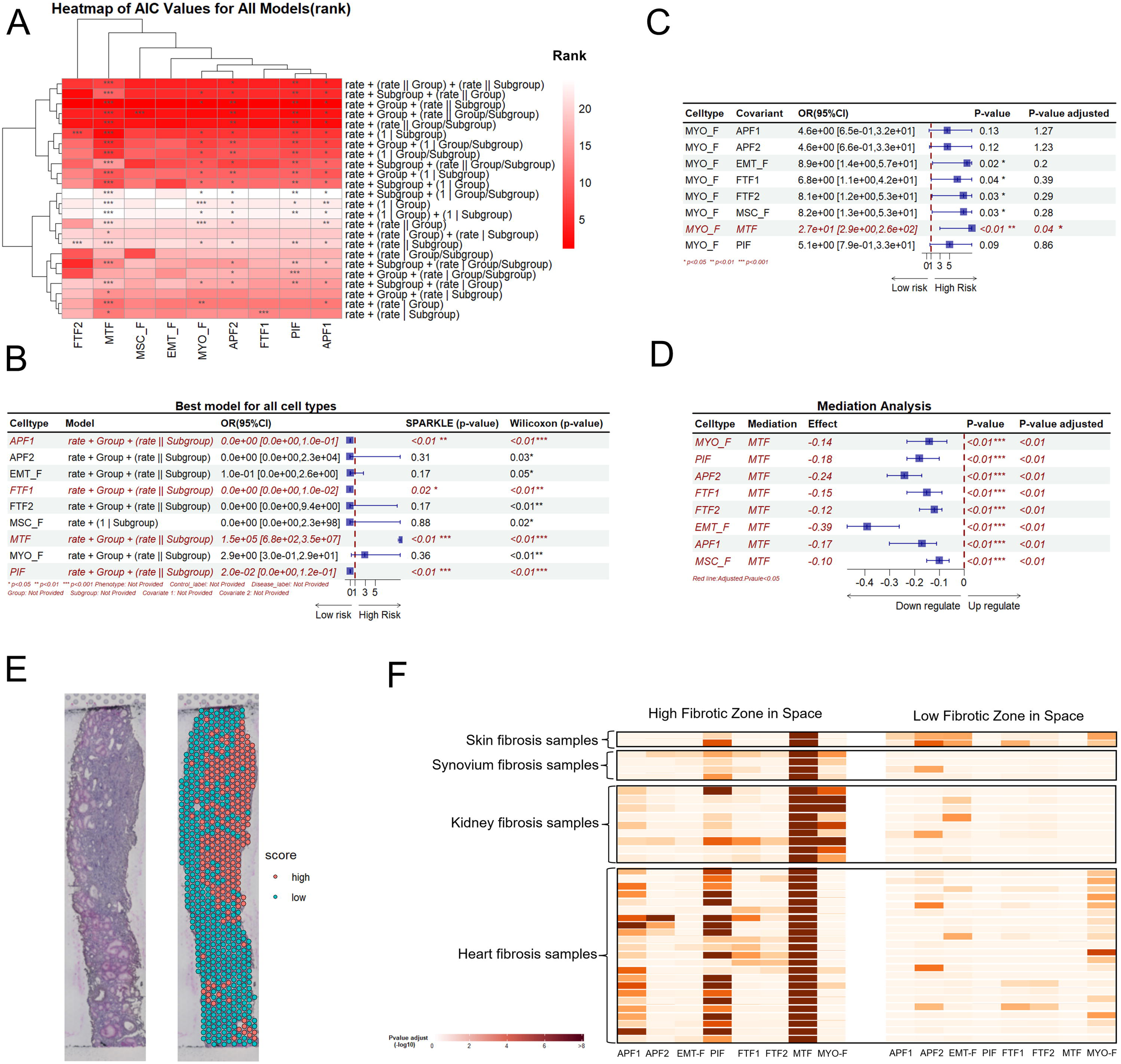
Exploration of the cell type–phenotype association with SPARKLE in FDFA. (A) Heatmap showing the AIC values and p values for all models across each cell type. The tissue information is included as a group, and the data source information is included as a subgroup. For each cell type, 24 logistic mixed effect models were used to calculate the AIC values and the p values of the rate coefficients for each model. The color represents the rank of the AIC values, with darker red indicating better models with lower AIC values. (B) Forest plot depicting the best model for each cell type–phenotype association. The model with the lowest AIC value for each cell type was selected as the best-fit model. Statistically significant associations with p values < 0.05 are colored red and italicized. The forest plot illustrates the odds ratio (OR) and its 95% confidence interval (CI). (C) Forest plot showing the best model when different cell ratios are added as covariates in the association analysis between phenotype and MYO-F. This finding suggests that MYO-F may be a risk factor that is independent of MTF. This phenomenon of changing conclusions when the cell type was introduced as a confounding factor was common and was observed in all datasets. This phenomenon is often easily overlooked in canonical cell type–phenotype analyses, resulting in erroneous conclusions. (D) Forest plot of the mediation analysis model with MTF in cell type– phenotype associations. (E) Representative images of H&E and MIA analysis of spatial transcriptome (ST) data. Each ST sample space is divided into low-fibrotic regions (green) and high-fibrotic regions (red). (F) Heatmap of the MIA analysis results. The colors indicate the proportions of different cell types in specific regions, with darker red representing a greater proportion of cells.

In fact, these masking and indirect mediation effects are widespread, as observed in all three other publicly available datasets (Supplementary Fig. 5D, 6D, and 7D; see Supplementary Fig. 5C, 6C, and 7C). SPARKLE effectively identifies the presence of mediating effects, aiding in the understanding of cell involvement in phenotype association analyses (Supplementary Fig. 5E, 6E, and 7E).

To validate our findings from the SPARKLE analysis, we used spatial transcriptomics data and multimodal intersection (MIA) analysis^22^ (Fig. 3E and 3F). A total of 38 samples were included in this analysis, including 2 samples of skin fibrosis (skin scar tissue)^23^, 4 of joint synovial fibrosis (rheumatoid arthritis)^24^, 9 of renal fibrosis (late-stage diabetic nephropathy)^25^, and 23 of myocardial fibrosis (postmyocardial infarction scar)^26^. Compared with that in the control region (low-fibrotic zone), the proportion of MTF significantly increased in the fibrotic region (high-fibrotic zone) (Fig. 3F). MYO-F also showed varying degrees of proportional increase in the fibrotic region.

Biologically, cell types often have extensive regulatory effects on each other. Moderation analysis of SPARKLE revealed that MYO-F positively regulated MTF, and MTF also had a statistically significant positive regulatory effect on MYO-F (Fig. 4A-4C). These results suggest that as MTF increases, its effects not only directly influence the phenotype but also amplify the promoting effects of MYO-F on the fibrotic phenotype through synergistic actions, further suggesting that MTF is a key cell type in the pathological process of multitissue fibrosis.

**Figure 4.**
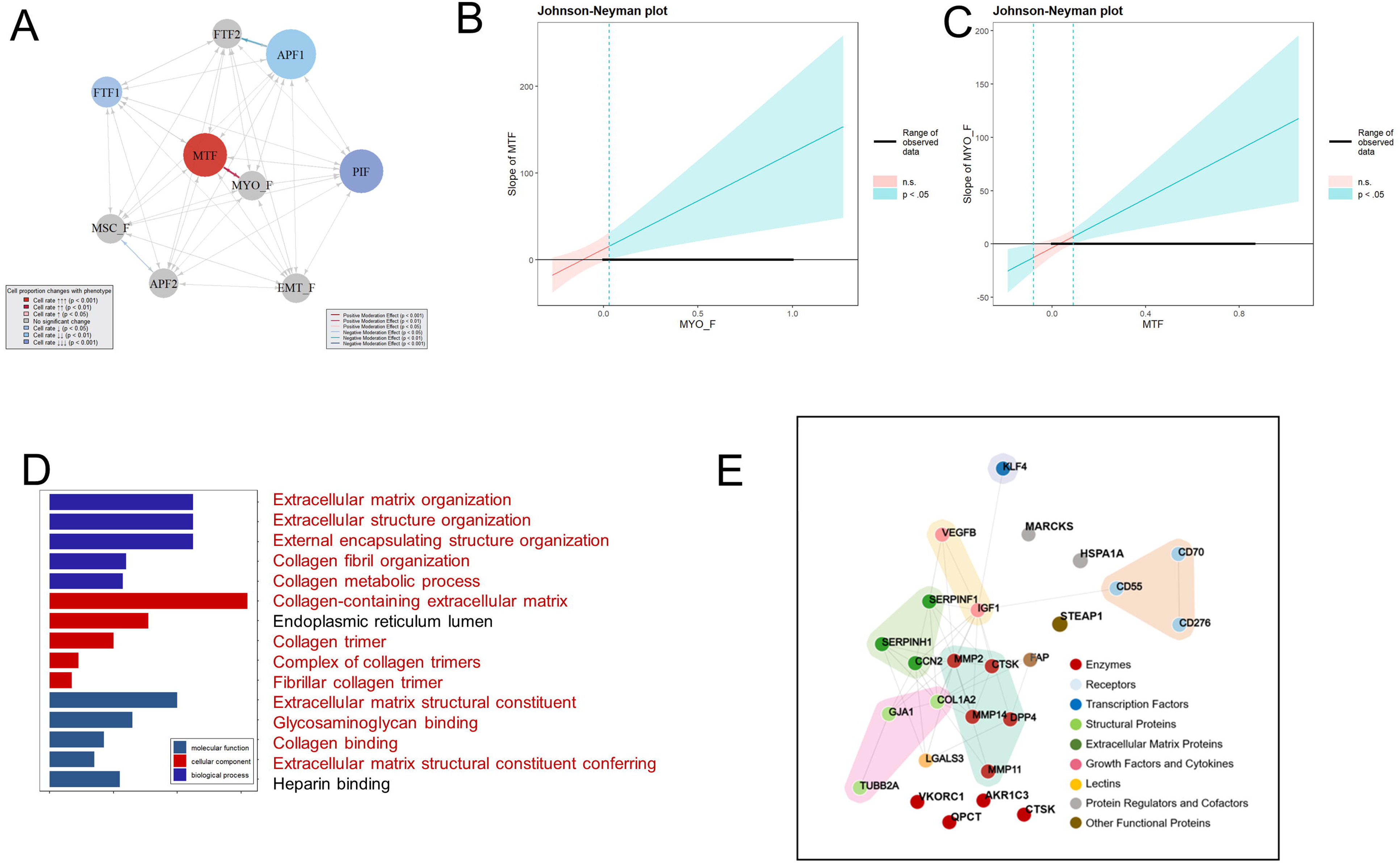
Moderation Analysis and Therapeutic Target Identification. (A) Network plot of the moderation analysis. The color of the spots represents the cell type–phenotype association, with red indicating a positive association and blue indicating a negative association. The arrows represent statistically significant moderation effects between cells and cell type–phenotype associations. Red arrows indicate a positive moderation effect, whereas blue arrows indicate a negative moderation effect. (B) Johnson–Neyman plots indicating the size and significance of the slope of the MTF proportion across all observed levels of MYO-F proportions. Shaded regions indicate 95% confidence intervals. (C) Johnson–Neyman plots indicating the size and significance of the slope of the MYO-F proportions across all the observed proportions of MTF. (D) Gene ontology analysis of potential fibrosis target genes (PFTGs). Fibrosis-related terms are colored in red. (E) Network plot of well-known druggable targets from PFTG. Different colors of points represent different target categories, and lines between points represent target interactions.

Furthermore, we attempted to identify MTF-based antifibrotic drug therapeutic targets^27^ (Fig. 4D,4E, also see Supplementary Table 3 for the full list). A total of 193 potential fibrosis target genes (PFTGs) were identified based on the significantly highly expressed marker genes of MTF in fibrosis-related diseases (p<0.01) (Supplementary Table 3).

## Discussion

By systematically curating published single-cell datasets, we created the cross-tissue, multidisease fibrotic disease fibroblast atlas (FDFA). With single-cell and spatial transcriptomic data from 11 common fibrotic diseases across six major human organs, FDFA is thus far the largest and most comprehensive single-cell atlas for fibrotic diseases. Because of its broad spectrum, we were able to identify 9 fibroblast subpopulations with distinct molecular profiles, providing a unique opportunity to identify cell type–phenotype associations in fibrosis globally.

Heterogeneity is extensive in single-cell datasets and is critical to understand. From an epidemiological perspective, the high heterogeneity of single-cell data primarily arises from two major types of bias: (1) selection bias and (2) information bias. First, selection bias occurs during sample collection, where specific population structures and sampling methods may lead to over- or underrepresentation of certain cell types, thereby affecting the generalizability of the results. Second, information bias stems from differences in sequencing platforms and data analysis techniques, which may lead to inconsistencies in data interpretation. The presence of these biases underscores the necessity of employing rigorous methods to control and correct for potential biases when analyzing single-cell data to ensure the accuracy and robustness of the results. Together, these sources of bias contribute significantly to the high heterogeneity observed in single-cell datasets, underscoring the importance of carefully addressing these variations to ensure accurate insights and reliable conclusions.

In particular, we found that, even after comprehensive data integration and batch correction, the impact of data source heterogeneity on cell proportions was comparable and even exceeded the effects of phenotypes. Canonical methods of cell type and phenotype association analysis in highly heterogeneous datasets can significantly impact the robustness of the results. With this knowledge, we developed the single-cell phenotype association research kit for large-scale dataset exploration (SPARKLE), a tool based on generalized linear mixed models (GLMMs) for large-scale single-cell cell–phenotype association analysis. SPARKLE supports the inclusion of multiple covariate variables as covariates to mitigate the impact of heterogeneity on result accuracy. Compared with traditional GLM, the GLMM-based approach of SPARKLE supports random effect models, improving estimation efficiency and preserving degrees of freedom. In fact, GLMM has been widely used in biological research, particularly in genome-wide association studies (GWASs), to account for population structure and relatedness, increasing the robustness of the results^42–44^. The possibility of applying the well-established methodology for a genetic-based association study to use in a cell type-based study is noteworthy. Furthermore, the well-established methodologies used in genome-wide association studies (GWASs) could also be applied to cell type-wide association studies (CWASs), suggesting promising potential for cross-disciplinary applications (Supplementary Fig. S8).

Based on SPARKLE, we identified MTF, which is significantly correlated with fibrosis, as confirmed through spatial transcriptomic MIA analysis. We found that MTFs contribute to fibrosis in direct and indirect ways. First, the proportion of MTFs significantly increases in fibrotic diseases, where MTFs highly express collagen-related genes and play crucial roles in multitissue fibrosis. Second, MTFs strongly interact with MYO-F, which synergizes with MYO-F, amplifying the impact of MYO-F on fibrotic phenotypes; MYO-F is a key effector cell in various fibrotic diseases^1,7,45^. Notably, the association between MYO-F and fibrotic phenotypes was masked by MTF, explaining why MYO-F did not show a significant association in isolation but became significant when controlling for MTF and other confounding cells.

In summary, SPARKLE introduces a novel paradigm for large-scale single-cell data analysis, addressing data heterogeneity and enhancing the accuracy of cell type– phenotype associations. Our findings identify matrix fibroblasts (MTFs) as a key subtype of fibrosis, offering new opportunities for broad-spectrum antifibrotic therapies. Additionally, the ability of SPARKLE to control for confounders and perform mediation and moderation analyses reveals deeper cell type–phenotype relationships, advancing our understanding of fibrotic diseases. By incorporating covariates, SPARKLE improves the accuracy and robustness of cell type–phenotype association analyses, making it a valuable tool for future research in complex diseases.

## Methods

### Cell–phenotype association analysis

The basic principle of cell type–phenotype association analysis in SPARKLE is based on a generalized linear mixed model (GLMM). The main form of the GLMM is as follows:

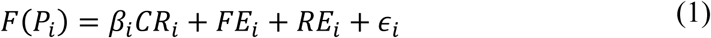

Here, *F*(∗) is the link function. SPARKLE supports various forms of link functions, with the default being the logit function, 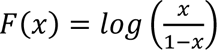.

*P_i_* is the phenotype of the *i*-*th* sample, which is a categorical variable. *CRi* is the proportion of cells in the *i-th* sample relative to the total. *βi* is the coefficient we want to calculate, representing the association between the phenotype and the cell proportion.

ϵ*i* is the random error for the **i*-th* sample.

The fixed effect part (*FE_i_*) is formulated as:

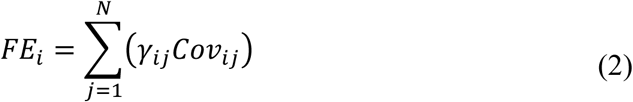

where *Covij* is the value of the *j*-th covariate for the fixed effect in the *i*-th sample. SPARKLE allows users to freely define multiple covariates for fixed effects. These covariates, considered confounding factors, are included in the fixed effects. The covariates can include information such as age and sex or proportions of other cell type ratios (*CR_i_*). The random effect part (*RE*) is also supported by SPARKLE and includes various random effect covariates. In accordance with previous studies^46^, the main forms of *RE* include the nest design, the nest and cross design, and the crossed and nested design. The *β*_*i*_ of the GLMM models are computed via the R package Lme4 (version 1.1.13)^47^. The heatmaps for visualization were created via pheatmap (version 1.0.12)^48^, and the forest plots were generated via the forestploter package (version 1.1.2)^49^.

### Hausman Test

In SPARLKE, the Hausman test is applied to determine whether the covariates should be random effects (*RE*) or fixed effects (*FE*)^50^. The Hausman test statistic *d* is calculated as follows:

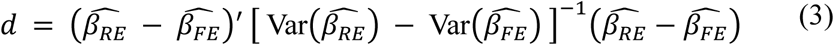

Here, 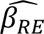 is the vector of coefficients estimated via random effects. 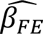 is the vector of coefficients estimated via fixed effects. 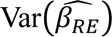 and 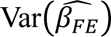 are the covariance matrices of the random effects and fixed effects estimates, respectively.

### Leave-one-out (LOO) test

The LOO test involves excluding one data source at a time and conducting the analysis. First, the total number of data sources in the integrated dataset is determined. Then, each data source is excluded sequentially, the remaining data are integrated, and the associations with the phenotype are calculated. The p values and significance (p < 0.05 was considered statistically significant) of the regression coefficients between different cell types and phenotypes were compared. Finally, the performances of different cell types across different data sources were compared.

### Bootstrap test

For each test dataset, we conducted proportional downsampling of each cell type to multiple levels: specifically, downsampling was performed to 90%, 80%, 70%, and so on, down to 10% of the original cell count. Following each downsampling step, we resampled to restore the original sample size. This entire procedure was repeated 1,000 times. After each iteration, we assessed the consistency of the cell type– phenotype association results to determine whether the associations remained stable across various downsampling levels.

### Cell–phenotype mediation analysis

Based on SPARKLE, mediation effect analysis between cell types and phenotypes can be conducted on the basis of GLMM. The basic principle is as follows:

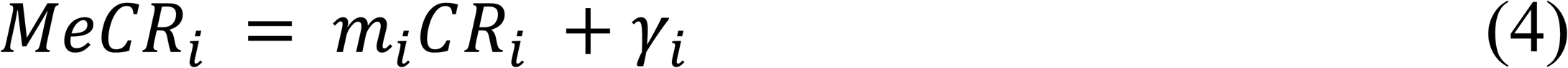

where *MeCR* is the proportion of the mediating cell type in the *i*-th sample. *CRi* is the proportion of the target cell type in the *i*-th sample. According to Equation (1), when considering *MeCR* as a covariate for the fixed effect, we have:

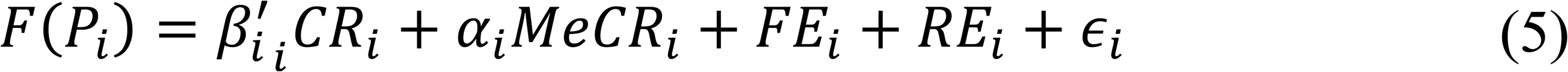

Substituting Equation (3) into Equation (4), we obtain:

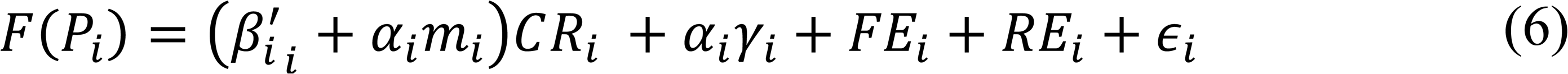

Here, 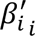 is the direct effect, and *αimi* is the indirect effect. By testing whether *αimi* is significantly different from zero, we can determine the presence of a mediating effect. If *αimi* and 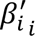 have the same sign and 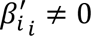, it indicates an indirect mediation effect. If *αimi* and 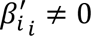 have the same sign and 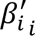, it indicates a complete mediation effect. If *αimi* and [inlie] have opposite signs, it indicates a masking effect. The mediation analysis in SPARKLE is performed mainly via the lme4 and bruceR packages (version 2023.9)^51^.

### Cell–phenotype moderation analysis

When the associations between the target cell and phenotype are studied, another cell type may have a moderating effect on the target cell. The calculation formula is as follows:

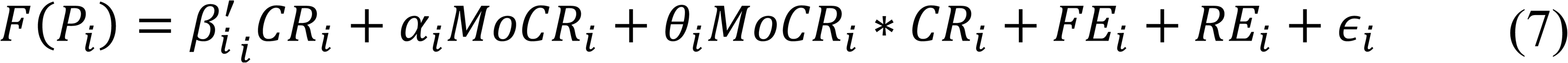

where *MoCRi* is the proportion of the cell type exerting a moderation effect on the *i*-th sample. If *θi* is significantly different from zero, the moderation cell ratio has a statistically significant moderating effect on the target cell. The moderation analysis in SPARKLE is also performed via the lme4 and bruceR packages.

### Data Acquisition

To ensure a precise search and selection of scRNA-seq data within the Gene Expression Omnibus (GEO) database, we downloaded all sample SOFT files from GEO^52^. SOFT files, a common data format in GEO, contain comprehensive details, including experimental design, sample information, raw data, and analysis results. By employing a keyword matching strategy within the SOFT files, we efficiently identified GSE entries in the GEO database that included single-cell sequencing data. We then downloaded the raw data corresponding to the identified GSE entries for subsequent data collection and processing.

### Data Processing

After downloading the data, we organized the expression matrices. We filtered the downloaded data by file type, processing those identified as potential single-cell expression matrices or metadata formats. Automatic searches within these files for gene and cell names were conducted. Files with row or column names matching gene or cell names were considered expression matrices. Data with too few cells or non-single-cell expression data were further filtered out. During this process, we also identified the gene version used in the expression matrix and the positions of gene and cell names within the files. All expression matrices were standardized and stored as h5ad files in the AnnData format.

### Metadata Collection

We subsequently collected metadata for the samples. For GEO data from which expression matrices could be extracted, we targeted the collection of phenotypic data. Metadata were extracted from the SOFT files corresponding to the samples provided by the submitters of the single-cell data. These files include the research background and sample descriptions. Although SOFT files follow certain structural rules, extracting phenotypic data from them is challenging. To increase annotation efficiency, we employed a large language model for annotation and validated the model’s accuracy by manually reviewing a subset of samples. The samples were annotated across multiple aspects, such as disease information, tissue or organ origin, treatment status, enrichment of specific cell types, and whether they were primary cultured cells. Disease information and tissue or organ data were further mapped to corresponding ontologies to provide structured annotations. Each dataset was also tagged with data quality metrics for each cell and dataset as outlined in the ACA^53^.

### Single-cell transcriptomic data processing

For all single-cell transcriptomics data in the FDFA, the count matrix was imported and processed with the Seurat (Version 5.1.0) package^54^. To prevent any interference in the analysis, potential doublets were predicted by Scrublet^55^. The merge function was employed to combine all individual objects into a single aggregate object, and the RenameCells function was utilized to ensure that all the cell labels were unique. Quality control was then performed based on several criteria: cells with fewer than 200 detected genes or more than 20% mitochondrial content were excluded. Additionally, cells with more than 6000 detected genes were removed to eliminate potential doublets. After filtering, 2,020,960 high-quality cells were retained for further analysis.

We then used the Harmony algorithm for data integration and batch effect correction^56^. A total of 147,254 fibroblasts were then identified from the integrated datasets. Clustering analysis was conducted based on edge weights between cells, creating a shared nearest-neighbor graph via the Louvain algorithm, implemented in the FindNeighbors and FindClusters functions. The resulting clusters were visualized via the UMAP method. For subclustering analysis, a similar procedure was followed, including normalization, selection of variably expressed features, dimensionality reduction, batch correction with Harmony, and cluster identification. Cell clusters were annotated by identifying differentially expressed markers via the FindAllMarkers function, which employs the default nonparametric Wilcoxon rank sum test with Bonferroni correction.

### FDFA Construction

Based on the previously obtained single-cell datasets, we performed screening and quality control. Initially, we removed 2,940 samples with incomplete information, resulting in 31,649 samples. Among these, 15,848 nonhuman samples were excluded, leaving 15,801 human single-cell samples.

Automated annotation was accomplished via ScType ^57^. We identified and excluded 11,510 samples that did not contain fibroblasts, retaining 4,291 samples with fibroblasts, totaling approximately 1.9 million fibroblasts.

We then screened the metadata to exclude nonfibrosis-related diseases while retaining normal fetal and adult samples as controls. Additionally, we included 38 spatial transcriptomic datasets of fibrotic disease samples.

Ultimately, we obtained 432 samples for the fibrotic disease fibroblast atlas (FDFA), comprising 394 single-cell transcriptomic samples and 38 spatial transcriptomic samples. The detailed information is listed in Supplementary Table 1.

### Cell functional analysis and gene enrichment analysis

The functional analysis of the cells was performed via the AddModuleScore function in the Seurat package. Functional gene sets were downloaded from MSigDB (https://www.gsea-msigdb.org/gsea/msigdb/). Gene enrichment analysis, including gene ontology (GO) and KEGG analysis, was conducted via the ClusterProfiler package (version 4.0)^58^. The top ten enriched GO terms and KEGG pathways ranked by the p value are depicted by the ggplot2 package. Protein–protein interaction information was downloaded from the STING database (version 12.0) ^59^. A network plot was generated with NORMA (https://pavlopoulos-lab-services.org/shiny/app/norma).

### Spatial transcriptomics data processing

We included 38 samples across four different fibrotic diseases (Supplementary Table 1). The Read10X_h5 and CreateSeuratObject functions from the Seurat package were used to create an object from the Space Ranger output. Using the Read10X_Image function, we loaded the H&E image data and normalized the dataset with the standard logNormalize function. The SpatialFeaturePlot function was used to display the expression levels of individual genes at their spatial locations. We subsequently combined our spatial transcriptomics (ST) data with integrated scRNA-seq datasets via the multimodal intersection analysis (MIA) method according to a previous study^22^. To determine the significance of the overlap between cell type marker genes and ST genes, we employed the hypergeometric cumulative distribution, using the entire gene set as the background for P value calculation. We divided the STs into highly fibrotic regions and less fibrotic regions. Different cell proportions in these regions were calculated separately.

### Epidemiology study design and participants

We utilized data from the National Health and Nutrition Examination Survey dataset of the United States (NHANES). This was a large cross-sectional study conducted by the National Center for Health Statistics to evaluate the overall health and nutritional status of the population in the U.S. This study was conducted in accordance with the guidelines approved by the Research Ethics Review Board of the National Center for Health Statistics (Protocol #2011-17, #2018-01). All patients involved in this study provided informed consent. We analyzed 301,258 individuals from the NHANES 2013-2020 database. Fibrotic disease diagnosis information and drug use information were documented in self-reported personal interview data on a variety of health conditions. The odds ratio (OR) was calculated via R software (version 4.2.2) in accordance with CDC guidelines (https://wwwn.cdc.gov/nchs/nhanes/tutorials). All of the estimates were calculated with sample weights to produce representative data of the civilian noninstitutionalized U.S. population.

## Supporting information

Suppplementary table 1

Suppplementary table 2

Suppplementary table 3

## Acknowledgments

The authors would like to thank Dr. Jing Zhang (Shanghai Tongren Hospital) for his work on the NHANES database and the drawing tools provided by Figdraw. We thank those who contributed to the NHANES data, including all anonymous participants in the study. This work was supported by funds from the National Key Research and Development Program of China (2022ZD0115004), as well as the State Key Laboratory of Protein and Plant Gene Research, the Beijing Advanced Innovation Center for Genomics (ICG) at Peking University, the Changping Laboratory, and the Shaw Foundation Hong Kong Limited. Part of the analysis was performed on the Computing Platform of the Center for Life Sciences of Peking University and supported by the High-performance Computing Platform of Peking University.

## Authors’ contributions

G.G., Y.X.Z. and L.K. designed the study; X.C., Y.D., S.Q.Y. and L.W. contributed to the data collection; X.C. analyzed the data; X.C. wrote the manuscript; all the authors helped revise this manuscript and approved it before publication.

Conceptualization: G.G., Y.X.Z. and L.K. Methodology: G.G., Y.X.Z. and L.K.

Data collection: X.C., Y. D and S.Q. Y

Visualization: X.C.

Funding acquisition: G.G., Y.X.Z. and L.K.

Supervision: G.G., Y.X.Z. and L.K.

Writing – original draft: X.C.

Writing – review & editing: G.G., Y.X.Z. and L.K.

## Availability of data and materials

FDFA is publicly available at https://zenodo.org/records/14011564. The SPARKLE package can be downloaded at https://github.com/gao-lab/SPARKLE.

The survey data are publicly available on the NHANES website (www.cdc.gov/nchs/nhanes/).

## Figure legend

**Supplementary Figure 1.**
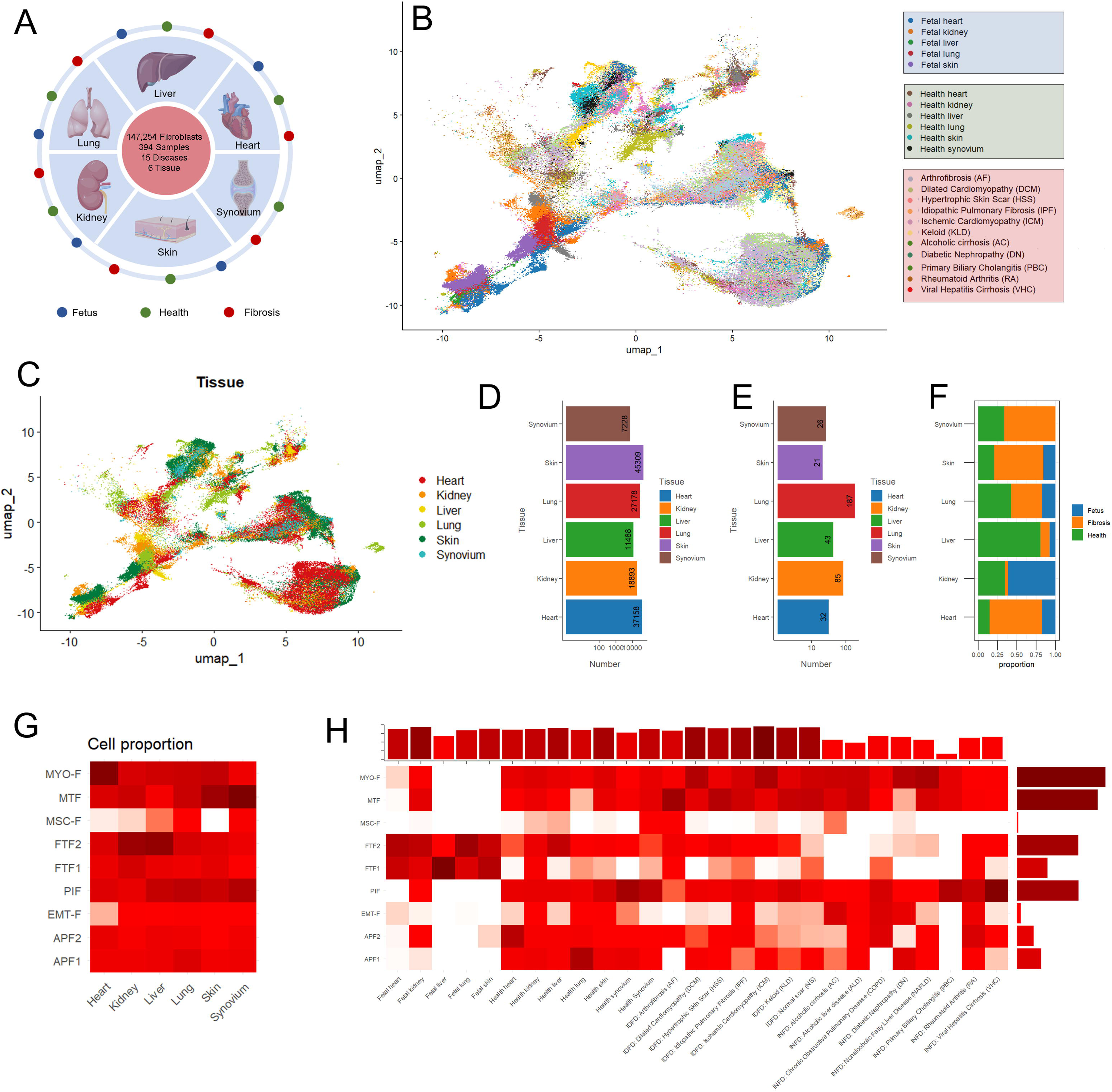
Analysis of the Fibrotic Disease Related Fibroblast Atlas (FDFA). (A) Schematic diagram of the data sources used in the FDFA. (B) UMAP visualization of fibroblasts across different diseases. (C) UMAP visualization of fibroblasts across different tissues. (D) Bar plot showing the cell counts in different tissues. (E) Bar plot showing sample counts in different tissues. (F) Stacked bar plot showing the proportions of cell counts from different sample sources across different tissues. (G) Heatmap showing the cell proportions of different fibroblast subtypes across different tissues. (H) Heatmap showing the cell proportions of different fibroblast subtypes across different diseases. The bar plot on the top of the heatmap depicts the cell counts in each disease, whereas the bar plot on the right side depicts the cell counts in each fibroblast subtype.

**Supplementary Figure 2.**
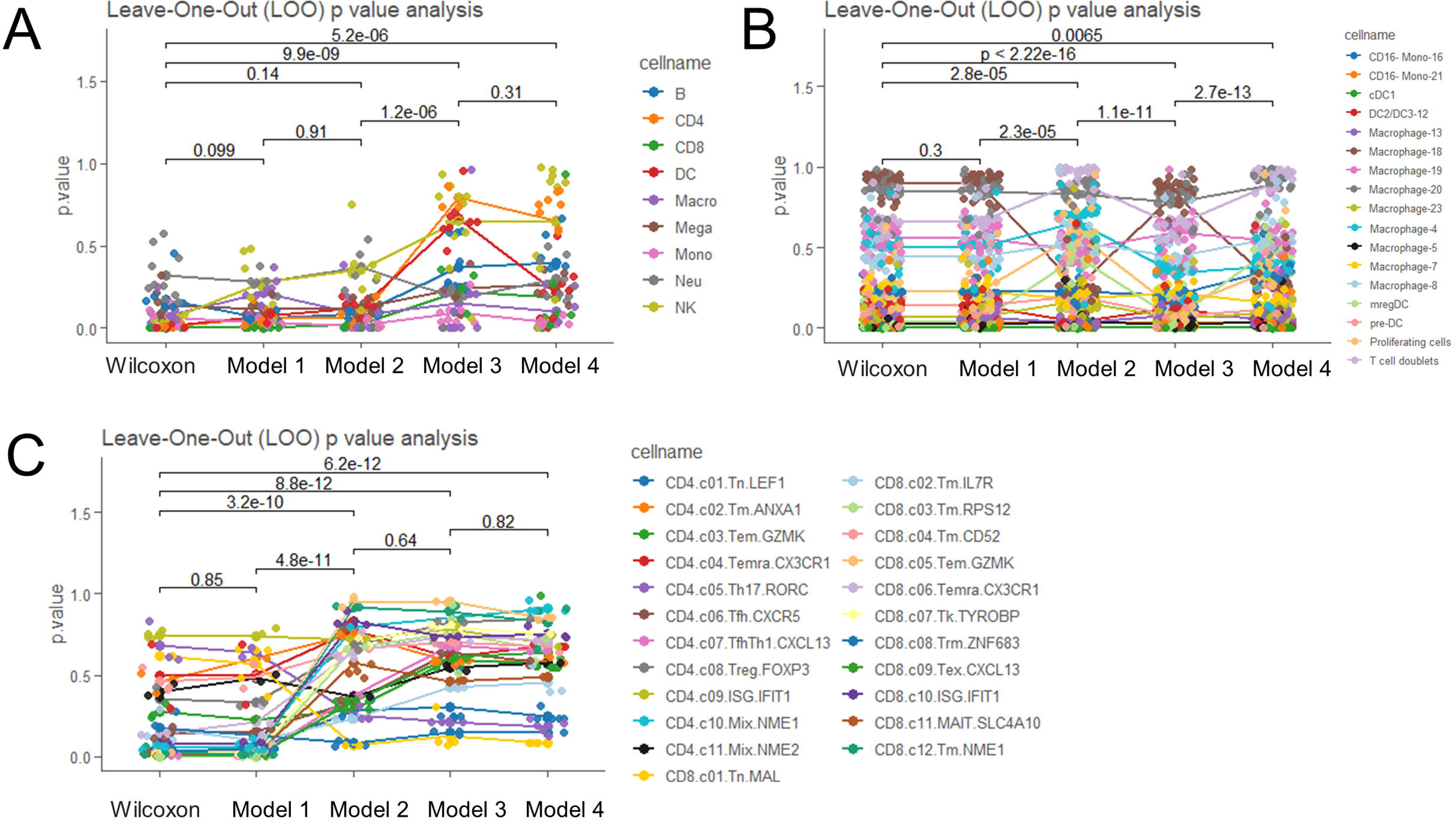
The Impact of Adding Covariate Information on Cell type–phenotype Association. These line charts present the effects of incorporating covariate information on the p values of cell type–phenotype associations across four different datasets: (A) the COVID-19 dataset, (B) the human monocyte and macrophage dataset, and (C) the tumor-infiltrating T-cell dataset. The five different model groups on the X-axis represent the following: Wilcoxon: Direct comparison of each cell type between two phenotype groups was conducted without using any covariate information. Model 1: Linear regression of the cell ratio and phenotype without any covariates was used to calculate the p value of the phenotype coefficient. Model 2: Linear regression with tissue information was added as a covariate to Model 1, calculating the p value of the phenotype coefficient. Model 3: Linear regression with data source information was added as a covariate to Model 1, calculating the p value of the phenotype coefficient. Model 4: Linear regression with both tissue and data source information was added as covariates to Model 1, calculating the p value of the phenotype coefficient. The Y-axis represents the p values of the associations between cells and phenotypes calculated by different models. Different colored points represent the p values of different cell types after leave-one-out (LOO) testing. The lines connect the mean p values for each cell type after LOO calculation. (P < 0.01, two-sided Wilcoxon test).

**Supplementary Figure 3.**
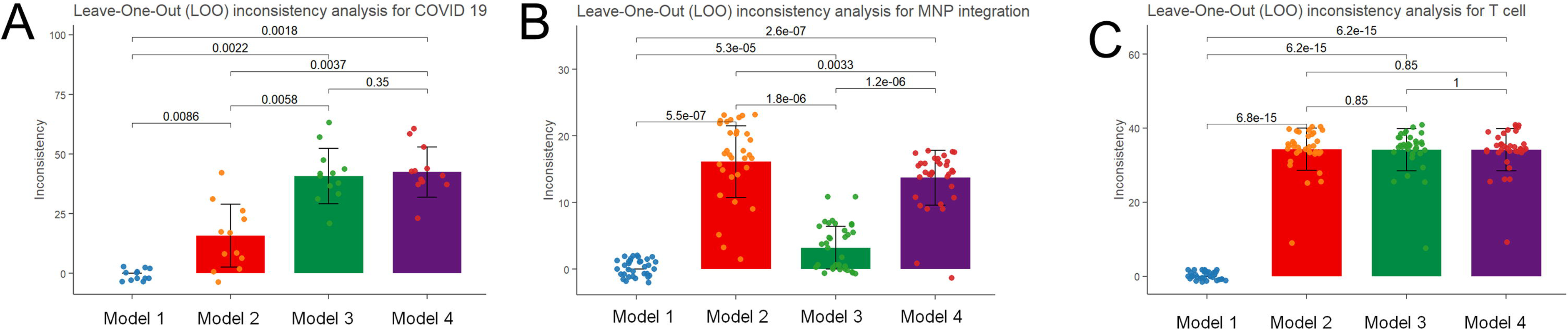
Comparison of Inconsistency Rates with Wilcoxon test Results after Adding Covariate Information. These bar plots present the inconsistency rates between the results of different models and the Wilcoxon test after adding covariate information across four different datasets: (A) the COVID-19 dataset, (B) the human monocyte and macrophage dataset, and (C) the tumor-infiltrating T-cell dataset. The four different model groups on the X-axis represent the following models. Model 1: Linear regression of the cell ratio and phenotype without using any covariate information, which was used to calculate whether the p value of the phenotype coefficient was statistically significant (p < 0.05). Model 2: Linear regression with tissue information was added as a covariate to Model 1, calculating the p value of the phenotype coefficient. Model 3: Linear regression with data source information was added as a covariate to Model 1, calculating the p value of the phenotype coefficient. Model 4: Linear regression with both tissue and data source information was added as covariates to Model 1, calculating the p value of the phenotype coefficient. The Y-axis represents the inconsistency rate, calculated as the proportion of cell types showing a change in the significance of their p values compared with the Wilcoxon results across all cell types. Different points represent inconsistent results after leave-one-out (LOO) testing. (P < 0.01, two-sided Wilcoxon test).

**Supplementary Figure 4.**
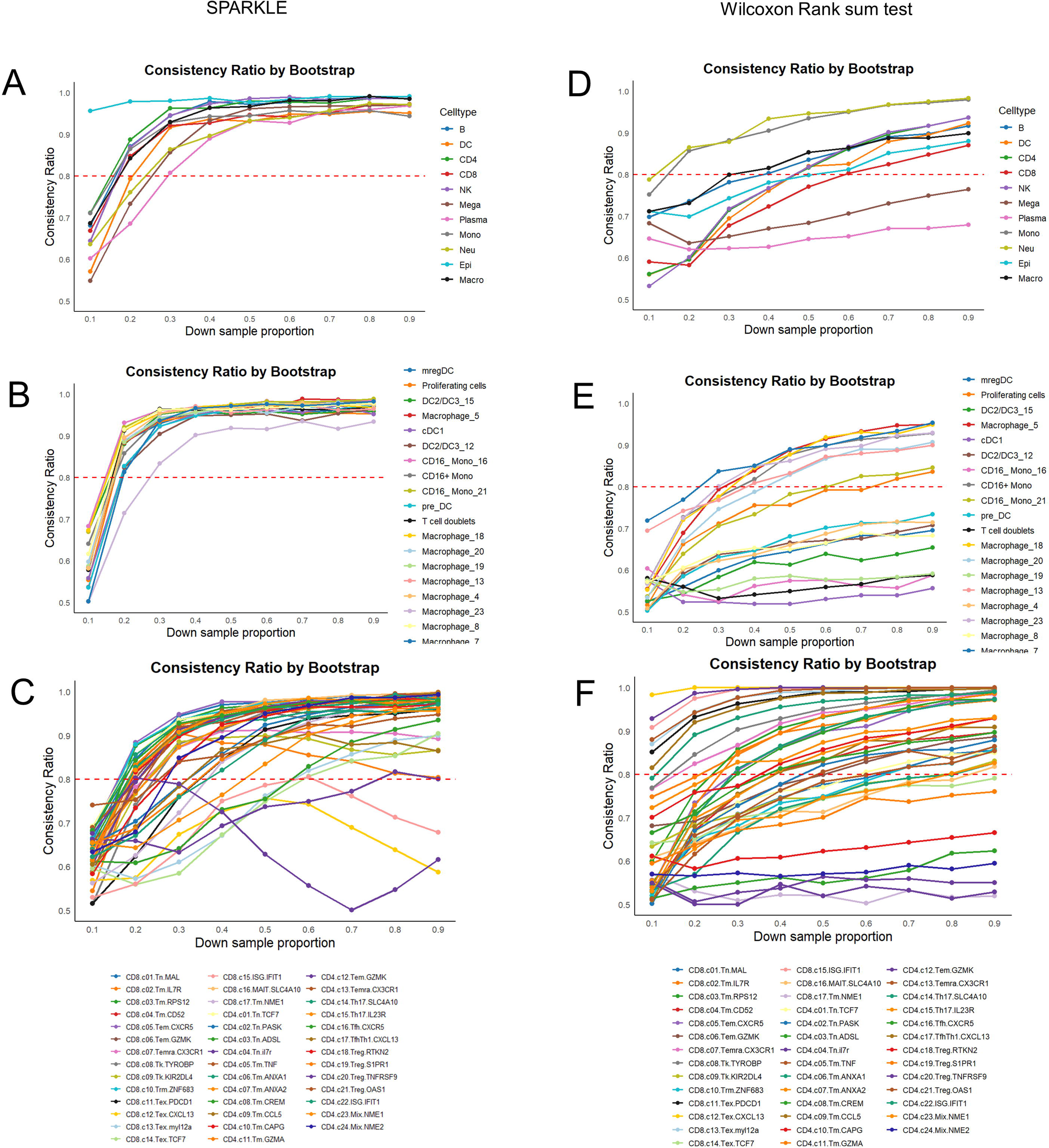
Bootstrap tests for SPARKLE and the Wilcoxon rank-sum test. This line plot illustrates the consistency ratios obtained from 1,000 bootstrap iterations. For each cell type in each dataset, samples were downsampled to a specific proportion and then upsampled to match the original sample size. Panels (A-C) show the SPARKLE consistency ratios during bootstrapping for the COVID-19 dataset (A), the human monocyte and macrophage dataset (B), and the tumor-infiltrating T-cell dataset (C). Panels (D-F) display the Wilcoxon rank-sum test consistency ratios for the same datasets: COVID-19 (D), human monocytes and macrophages (E), and tumor-infiltrating T cells (F). Statistical significance was assessed via two-sided Wilcoxon tests (P < 0.01).

**Supplementary Fig. 5.**
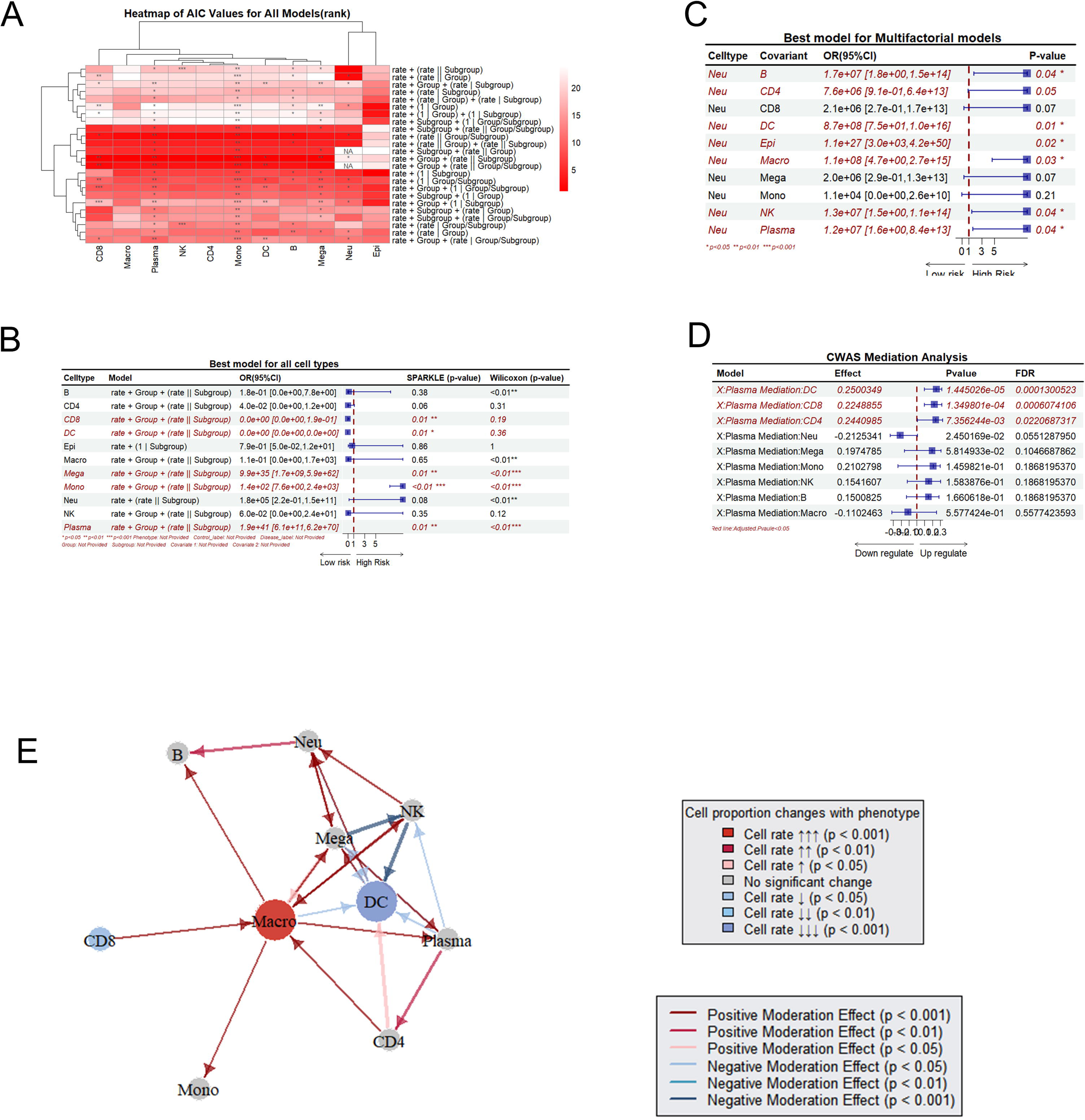
Exploration of the cell type–phenotype association with SPARKLE in the COVID-19 dataset. (A) Heatmap showing the AIC values and p values for all the models across each cell type. The color represents the rank of the AIC values, with darker red indicating better models with lower AIC values. (B) Forest plot depicting the best model for each cell type–phenotype association. Statistically significant associations with p values < 0.05 are colored red and italicized. The forest plot illustrates the odds ratio (OR) and its 95% confidence interval (CI). (C) Forest plot showing the best model when different cell ratios are added as covariates in the association analysis between phenotype and the mono/plasma ratio. (D) Forest plot of the mediation analysis model of the effects of plasma on cell type–phenotype associations. (E) Network plot of the moderation analysis. The color of the spots represents the cell type–phenotype association, with red indicating a positive association and blue indicating a negative association. The arrows represent statistically significant moderation effects between cells and cell type–phenotype associations. Red arrows indicate a positive moderation effect, whereas blue arrows indicate a negative moderation effect.

**Supplementary Fig. 6.**
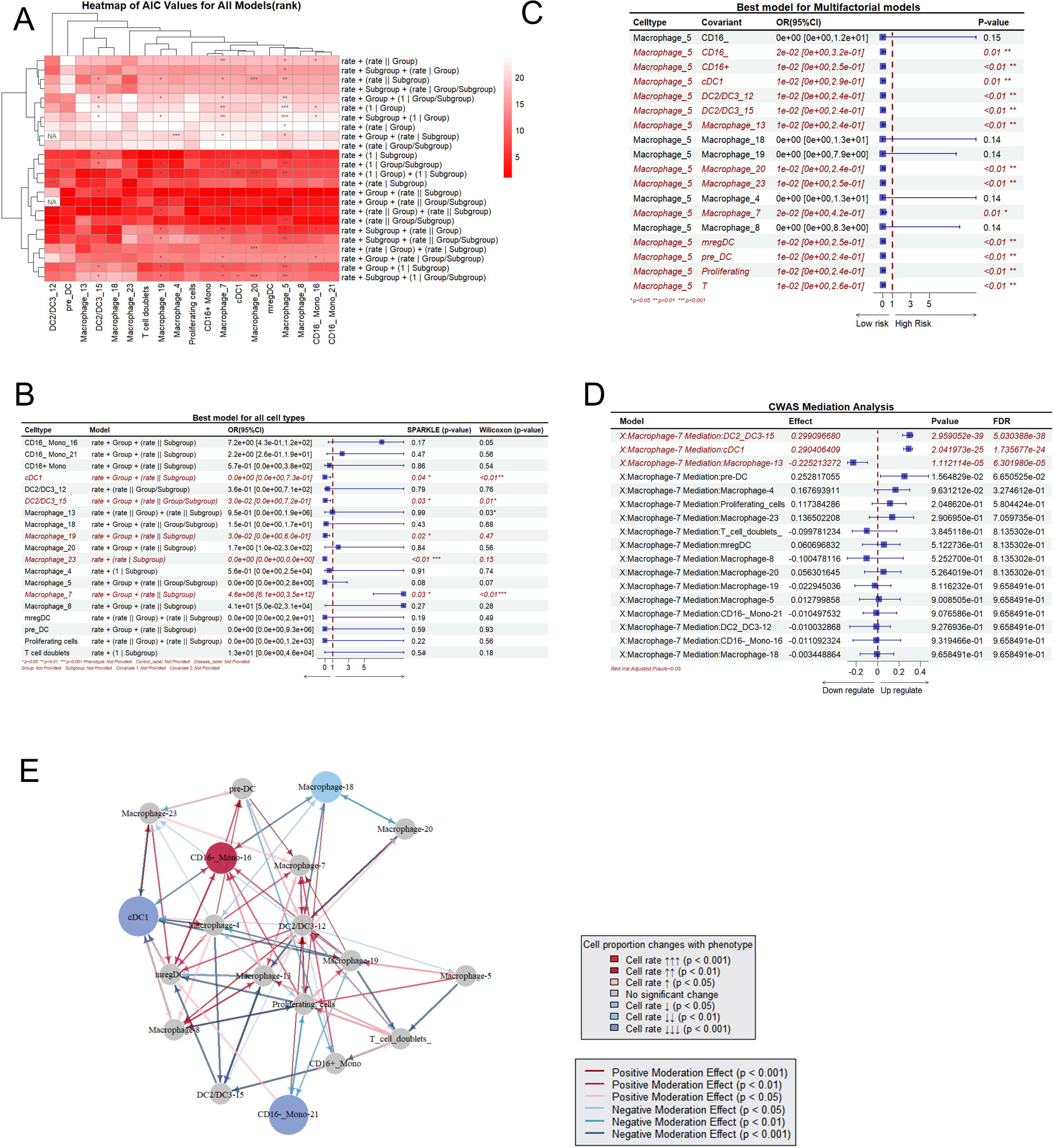
Exploration of the cell type–phenotype association with SPARKLE in the human monocyte and macrophage dataset. (A) Heatmap showing the AIC values and p values for all the models across each cell type. The color represents the rank of the AIC values, with darker red indicating better models with lower AIC values. (B) Forest plot depicting the best model for each cell type– phenotype association. Statistically significant associations with p values < 0.05 are colored red and italicized. The forest plot illustrates the odds ratio (OR) and its 95% confidence interval (CI). (C) Forest plot showing the best model when different cell ratios were added as covariates in the association analysis between phenotype and macrophages 5. (D) Forest plot of the mediation analysis model of the role of Macrophage7 in cell type–phenotype associations. (E) Network plot of the moderation analysis. The color of the spots represents the cell type–phenotype association, with red indicating a positive association and blue indicating a negative association. The arrows represent statistically significant moderation effects between cells and cell type–phenotype associations. Red arrows indicate a positive moderation effect, whereas blue arrows indicate a negative moderation effect.

**Supplementary Fig. 7.**
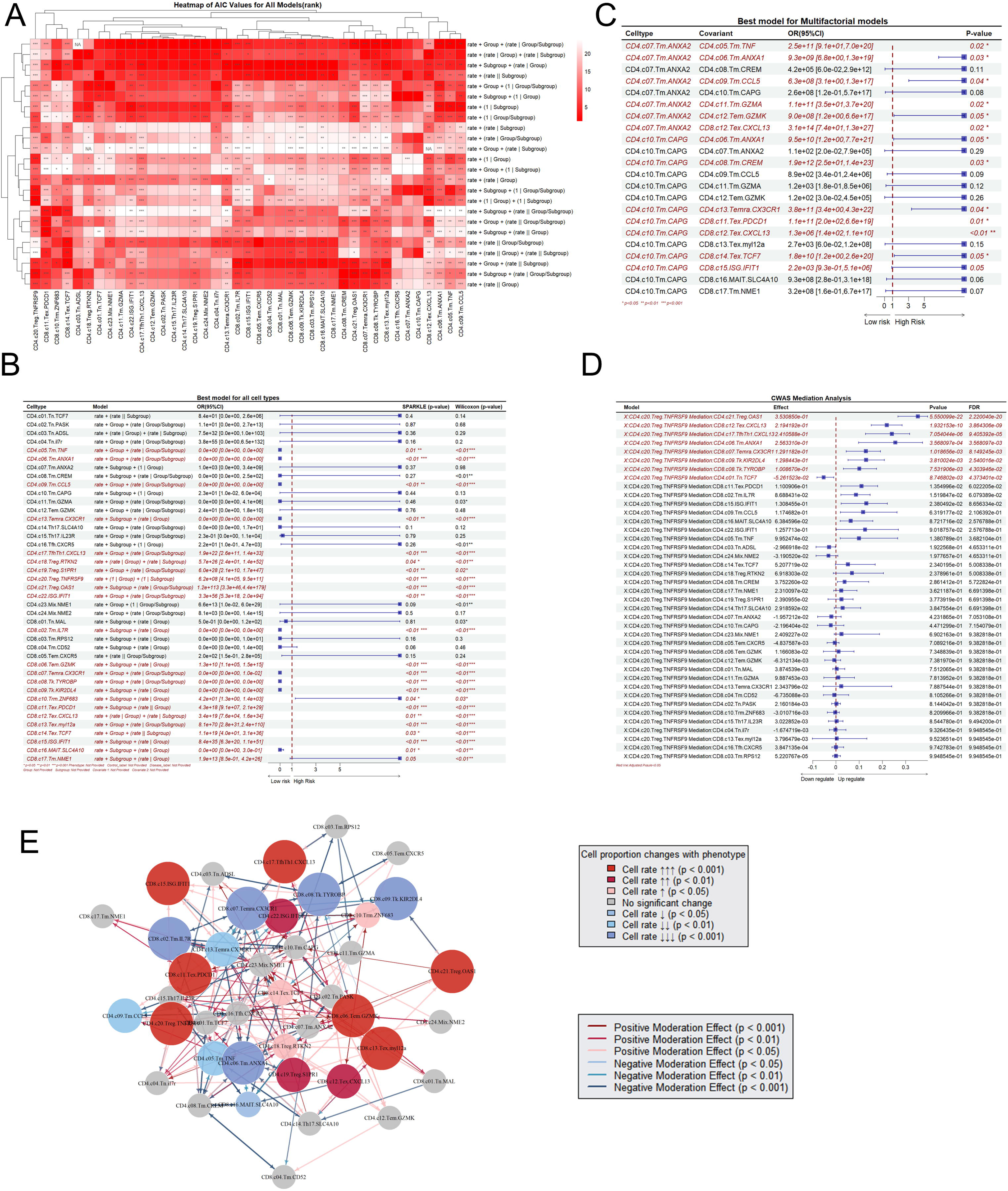
Exploration of the cell type–phenotype association with SPARKLE in the tumor-infiltrating T-cell dataset. (A) Heatmap showing the AIC values and p values for all the models across each cell type. The color represents the rank of the AIC values, with darker red indicating better models with lower AIC values. (B) Forest plot depicting the best model for each cell type–phenotype association. Significant associations with p values < 0.05 are colored red and italicized. The forest plot illustrates the odds ratio (OR) and its 95% confidence interval (CI). (C) Forest plot showing the best model when different cell ratios were added as covariates in the association analysis between phenotype and CD8.C06.Temra.CX3CR1. (D) Forest plot of the mediation analysis model of CD8.05.Tem. GZMK in cell type–phenotype associations. (E) Network plot of the moderation analysis. The color of the spots represents the cell type–phenotype association, with red indicating a positive association and blue indicating a negative association. The arrows represent statistically significant moderation effects between cells and cell type–phenotype associations. Red arrows indicate a positive moderation effect, whereas blue arrows indicate a negative moderation effect.

**Supplementary Fig. 8.**
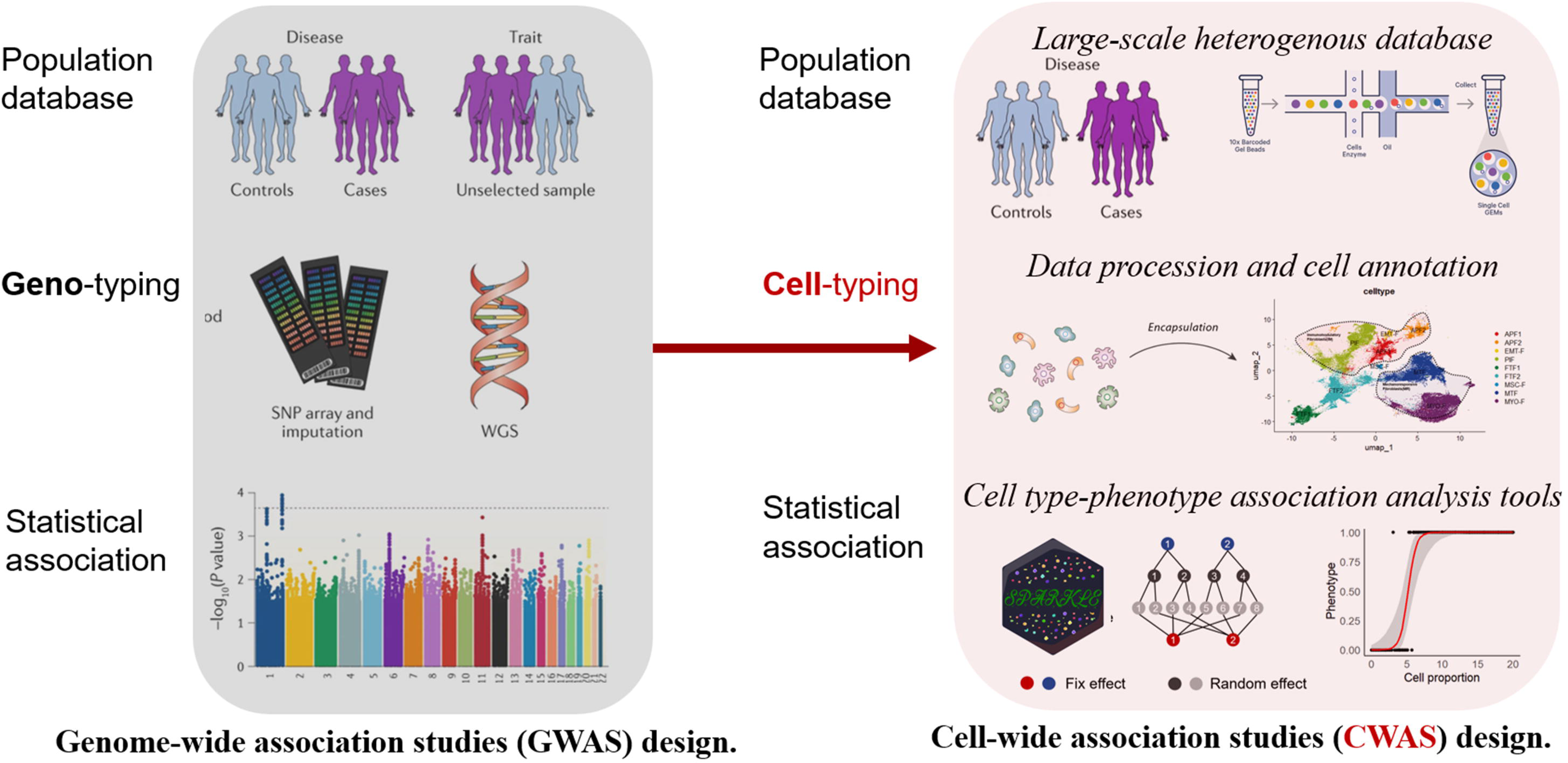
Comparison between Genome-Wide Association Studies (GWAS) and Cell-Wide Association Studies (CWAS). On the left, the traditional GWAS approach is illustrated. Population-based datasets consisting of controls, cases, or unselected samples are used for genotyping through methods such as SNP arrays and whole-genome sequencing (WGS). Statistical association tests are then applied to identify genetic variants associated with disease or traits. On the right, the CWAS framework, a methodology inspired by GWAS, is depicted. Analytical methods developed for GWASs can be adapted and applied to CWAS, enhancing the ability to identify associations between cell type proportions and phenotypes. CWAS provides a new perspective on phenotype associations, enabling insights that extend beyond the scope of traditional GWAS.

